# Potential for metal-coupled methane oxidation by *Candidatus* Methanocomedenaceae in coastal sediments

**DOI:** 10.64898/2026.03.20.712598

**Authors:** Anna J. Wallenius, Andy O. Leu, Robin Klomp, Simon J. McIlroy, Gene W. Tyson, Caroline P. Slomp, Mike S.M. Jetten

## Abstract

Anaerobic methanotrophic (ANME) archaea are important players in the microbial methane cycle, mitigating methane emissions from anoxic environments. ANME are found ubiquitously in methane-rich sediments, where they can couple anaerobic methane oxidation (AOM) to different electron acceptors such as sulfate, metal oxides, and natural organic matter (NOM). However, we still lack understanding of the geochemical niches and preferred metabolic pathways of most ANME subclades. Here, we investigated the genomic potential and ecophysiology of ANME-2a with respect to metal-dependent AOM in brackish metal-rich coastal sediments. We assembled several high-quality ANME MAGs from subclades with high strain heterogeneity and analyzed the genomic potential for metal-AOM. Additionally, we monitored long-term enrichments with various electron acceptors from the same sediments. Ultimately, we recovered 8 novel genomes of ANME-2a that clustered with an uncharacterized genus with only 2 representatives in public databases for which we propose the name ‘*Candidatus* Methanoborealis’. The analysis of the MAGs showed two different clusters within this genus; one comprising of MAGs from the Baltic Sea that showed high potential for extracellular electron transfer (EET) required for metal-AOM, and another cluster form more diverse environments with less EET potential. The Baltic Sea *Ca.* Methanoborealis were the only canonical methanotrophs in the incubations during active methane oxidation and metal reduction. Our results contribute to the understanding of the phylogenomic and metabolic diversity in ANME subclades, which will help to further characterize novel ANME lineages from complex sediment samples.

## Introduction

Anaerobic methane oxidizing archaea (ANME) are important players in microbial methane-cycling as they remove significant amounts of methane produced in anoxic sediments thereby limiting its escape into the overlying water column and atmosphere. In marine sediments, methanotrophic ANME couple anaerobic oxidation of methane (AOM) to sulfate reduction in a consortium with sulfate-reducing bacteria (SRB; Boetius et al., 2000). ANME use the reverse methanogenesis pathway for methane oxidation, and the electrons are transferred to SRB by interspecies electron transfer [2]. In deeper sulfate-depleted layers, as well as freshwater and brackish sediments, ANME can use alternative electron acceptors for AOM, such as, nitrate, metal oxides or humic acids [3–6]. However, the lack of highly enriched cultures still limits our ability to study the cellular mechanism driving AOM with different electron acceptors.

ANME archaea belong to three phylogenetically different clades within the phylum Halobacteriota (ANME-1, 2 and 3). ANME-1 (*Candidatus* Methanophagales) are the most distant related group, whereas ANME-2 and 3 belong to the Methanosarcinales order together with many methanogenic archaea [7]. ANME-2 form three separate families; *Ca.* Methanogasteraceae (ANME-2c) *Ca.* Methanoperedenaceae (ANME-2d) and *Ca*. Methanocomedenaceae that includes ANME-2a and 2b. The majority of physiological and metagenomic studies have focused on freshwater *Ca.* Methanoperedenaceae, that are more frequently sampled [5,8,9]. Species from this family have been shown to couple AOM to reduction of nitrate, iron and manganese oxides, and current generation without a syntrophic partner [5,4,10,11]. *Ca.* Methanoperedenaceae genomes encode several multiheme cytochrome proteins (MHC), which have been proposed to mediate extracellular electron transfer (EET) to external electron acceptors such as metal oxides, without the need of a bacterial partner [11,12].

*Ca*. Methanocomedenaceae (ANME-2ab) are commonly found in the sediments of marine and brackish ecosystems based on 16S rRNA gene amplicon sequencing (Aromokeye et al., 2020; Dalcin Martins et al., 2024; Rasigraf et al., 2020; Wallenius et al., 2025), but comprehensive studies on their physiology and differences between the subclades are still lacking as bioreactor or environmental studies do not often distinguish between the different subclades [18,19]. Metagenomic studies on marine sediments have distinguished several subclades phylogenomically within ANME-2ab; ANME-2b with only one described genus (*Ca*. Methanomarinus) and ANME-2a that is currently divided into three different genus level clades; *Ca.* Methanocomedens with 20 publicly available MAGs, “QBUR01” (15 MAGs) and “Kmv04” (2 MAGs; GTDB-Tk v2.4.1 Release 226; Chaumeil et al., 2022).

Based on batch incubations and amplicon and metagenomic sequencing studies, methane oxidation by ANME-2ab has been linked to reduction of iron (Fe-AOM) as well as natural organic matter (NOM-AOM) in brackish (iron-rich) sediments as well as sulfate-free mud volcanoes [13,15,20–22]. However, how ANME-2a mediate Fe-AOM or NOM-AOM and whether they require a syntrophic partner is not known. In marine/brackish sediments their abundance usually correlates quite well with that of *Desulfuromonadales* abundance, but in the presence of an artificial electron acceptor they have also been shown to be capable of AOM alone [23]. Long-term incubations with methane and ferrihydrite resulted in enrichment of ANME-2a and uncultured *Desulfobulbaceae* [24]. ANME-2a genomes encode multiple MHCs and a homologue for bacterial OmcZ, a protein linked to nanowire formation used in EET in *Geobacter* [7,25,26]. However, due to the low number of ANME-2ab cells in enrichments and the lack of representative metagenomically assembled genomes (MAGs), the mechanism of metal-AOM as well as their putative bacterial partners is not well explored.

In this study, we aimed to retrieve and characterize ANME-2a MAGs from different coastal systems where they are present [15,27] but have been difficult to obtain due to sample complexity, low abundance and high strain heterogeneity. We combined a range of assembly and binning strategies such as differential coverage binning to retrieve representative genomes to analyse their genomic potential for methane oxidation coupled to metal reduction [28]. Previously, we showed active metal-AOM in batch incubations from two sites in the Bothnian Sea {14-16;65], and used these cultures to establish long-term laboratory AOM incubations with iron, manganese and graphene oxide (humic acid analogue) as sole electron acceptor, to obtain a high relative abundance of ANME-2ab and thus be able to better obtain their genomic blueprint. All MAGs recovered are affiliated with *Methanocomedenaceae* ANME-2a [29]. We analyzed the MAGs of this novel group to unravel their potential AOM mechanisms, range of electron acceptors, and syntrophic partners.

## Materials and methods

### Sample origin

The samples obtained in this study originate from two coastal sites in the Bothnian Sea (BS), NB8 (33 m depth; 63°29.3’N, 19°46.2’E) and US2 (200 m depth; 62°50.99’ N; 18°53.53’ E), sampled in 2019 and 2022, respectively. The information on the sampling and processing of the original sediment samples is given in Supplementary methods. The US5B (Bothnian Sea) and GV (Lake Grevelingen) sediment samples originate from different studies [14-16;65], but information about the location, sampling, sample processing and MAG recovery is described in Supplementary methods.

### DNA extraction

For sediment and incubation samples, DNA was isolated with the DNeasy PowerSoil Pro DNA isolation kit (Qiagen, Venlo, Netherlands). For the incubation samples, in addition to the normal protocol, according to the kit’s trouble shooting guide, we added a pre-incubation of the samples at 65°C for 10 min with CD1 solution, to improve DNA recovery from cells that are hard to lyse such as archaeal cells. The lysed cells were bead-beaten on a TissueLyser LT (Qiagen) for 10 min at 50Hz and. The samples from graphene oxide amended samples were treated with 2% polyvinylpyrrolidone (PVP) before cell lysis to prevent DNA adsorption by graphene oxide.

### 16S rRNA gene amplicon sequencing

Archaeal and bacterial 16S rRNA amplicon sequencing was performed on the Illumina MiSeq Next Generation Sequencing platform by Macrogen (Seoul, South Korea) using Herculase II Fusion DNA Polymerase Nextera XT Index Kit V2, yielding 2x300bp paired end reads. For archaea we used primers Arch349F (5′-GYGCASCAGKCGMGAAW-3′) and Arch806R (5′-GGACTACVSGGGTATCTAAT-3′; Takai and Horikoshi, 2000) and for bacteria Bac341F (5′-CCTACGGGNGGCWGCAG-3′; [31] and Bac806R (5′-GGACTACHVGGGTWTCTAAT-3′; [32]. The obtained reads were analysed as described before [27]. Raw reads are accessible on the National Center for Biotechnology Information (NCBI) website under the accession number PRJNA1310174.

### Metagenomic sequencing

Metagenomic sequencing was performed with a TruSeq DNA PCR free library using an insert size of 550 bp on a NovaSeq6000 or NovaSeq X (Illumina) platform, producing 2 × 151 bp paired-end reads (10Gb per sample). The US2 sample and a sample from NB8 incubations with manganese were additionally sequenced with long read sequencing with Nanopore. Library preparation was done starting with 150 to 400 ng of DNA, quantified by Qubit. The quality of the DNA was checked by agarose gel electrophoresis. DNA Library construction was performed using the Native Barcoding Kit 24 V14 (SQK-NBD114-24), according to the manufacturer’s protocol (Oxford Nanopore Technologies, Oxford, UK). The library was loaded on a Flow Cell (R10.4.1) and sequenced using the MinION Mk1C device (Oxford Nanopore Technologies, Oxford, UK), according to the manufacturer’s instructions. Quality of the sequence reads was analyzed using fastqc [33].

SingleM v0.13.2 was applied to each metagenomic paired-end dataset to determine counts for discrete sequence-based operational taxonomic units (OTUs) using the ‘pipe command’ (https://github.com/wwood/singlem). Based on the coverage of each ANME-clade, we used low-abundance k-mer trimming using the khmer script trim-low-abund.py prior to differential coverage binning [28]. The trimming parameters depended on the coverage of ANME per sample. The filtered reads were assembled and binned using the Aviary pipeline (v.0.11.0; Newell et al., 2024). The reads were trimmed with fastp ( v0.24.0; Chen, 2023), assembled with metaSPAdes (v. 4.0.0; Nurk et al., 2017), mapped with CoverM ( v0.7.0; Aroney et al., 2025) and binned with Vamb (v3.0.2; Nissen et al., 2021), MetaBAT and MetaBAT2 (v2.15; Kang et al., 2015, 2019), Semibin [41], Rosella (v0.5.5; https://rhysnewell.github.io/rosella/) and DASTool (v. 1.1.2; Sieber et al., 2018). The metagenomically assembled genomes (MAGs) were taxonomically classified with GTDB-Tk release 220 (v2.4.0; Chaumeil et al., 2020). MAG completeness and contamination was estimated with CheckM2 (v1.0.2; [44]. Scaffolding and gap-filling of the metagenome assembly was performed using the “roundup” mode of FinishM v0.0.7 (https://github.com/wwood/finishm). The resulting ANME bins were further refined with MAGpurify (https://github.com/snayfach/MAGpurify). Genomes were annotated with DRAM v1.0 [45] and metascan [46]. Genes involved in metal reduction were searched for with FeGenie [47]. The ANI between the MAGs was determined with FastANI (v.1.33; Jain et al., 2018) and AAI was calculated with EzAAI [49]. Proteins were identified with prodigal ( v2.6.3; Hyatt et al., 2010) and used to search for proteins with heme-binding motifs (>3).

### Phylogenetic trees

*mcrA* sequences identified in this study were processed and aligned with published reference sequences [7] using mafft --auto and a maximum likelihood tree generated with IQ-TREE 2.1.4-beta [51] using flags -m MFP -B 1000 -nt AUTO. The tree was edited and annotated with iTol v7 [52]. For the genome-tree we used the multiple sequence alignment file generated with GTDB-Tk classify_wf function using 120 archaeal concatenated single-copy marker genes for a subset of known ANME and related species. The tree was generated and annotated as detailed above for the *mcrA* tree.

### Birnessite and ferrihydrite synthesis

Birnessite was synthesized as described before by reducing KMnO_4_ with C_3_H_5_NaO_3_ at ambient temperature and pressure [53]. Briefly, 5 mL of 60% C_3_H_5_NaO_3_ was mixed with 500 ml of 1 g L^-1^ KMnO_4_ for 2.5 h. The birnessite was harvested by centrifugation and washed five times with Milli-Q water and freeze-dried. Ferrihydrite was synthesized by adjusting pH of a 0.1 M Fe(NO_3_)_3_ solution to 7.5 with 1M KOH as described before [54] while vigorously stirring. The iron minerals were harvested by centrifugation and washed five times with Milli-Q water.

### Incubation setup

The NB8 and US2 incubations were started by mixing pre-incubated sediment ( see details in supplement) with sulfate-free artificial seawater, adding ^13^C-CH_4_ as the electron donor and either ferrihydrite, birnessite or graphene oxide (Sigma-Aldrich, Darmstadt, Germany) as the only electron acceptor. The NB8 enrichments of Mn-AOM and Fe-AOM were 780 and 345 days old, respectively and US2 Mn-AOM, Fe-AOM and GO-AOM 200 days old when 10 ml of each enrichment incubation was transferred to a new serum bottle and diluted with 50 ml of ASW. The ASW had a different salinity for NB8 (∼3.5) and US2 (∼6) and the compositions are listed in Supplementary methods. Headspace was replaced with N_2_ and amended with 20 % ^13^C-CH_4_ and 2 % CO_2_ to reach an overpressure of 1.5 bar. Ferrihydrite (7.5 mM), birnessite (5 mM) and graphene oxide (4 mg L^-1^) were added to the new slurry. The samples for NB8 were incubated at 4°C and US2 at room temperature (∼20°C) by gently shaking in the dark. Methane oxidation activity was tested every 2-3 months with GC-MS (Agilent 5977B System, Waldbronn, Germany) injecting 50 µl of headspace gas to determine the ^13/12^C ratio of headspace CO_2_. The samples were either newly diluted (1:5) or allowed to settle and 90% of the supernatant was replaced with new ASW and trace elements every 6-8 months, taking 2 ml of the slurry for DNA analysis. Samples for sulfate and dissolved metals were taken from the supernatant before and after sample dilution or change of ASW.

### Analytical procedures

The supernatant was filtered over 0.45 µm pore size filters and subsampled for the analysis of sulfate (SO_4_^2-^) and dissolved manganese (Mn^2+^) and iron (Fe^2+^). The samples for metals were acidified with 10 µL 35% suprapure HCl per milliliter of sample and stored at 4°C until analysis. Concentrations of SO_4_^2-^ were determined via ion chromatography (Metrohm 930 Compact IC Flex; detection limit of 10 µmol L^−1^). Mn^2+^ and Fe^2+^ were measured via ICP-OES (PerkinElmer Avio; detection limit 0.1 µmol L^−1^ for Fe, 0.02 µmol L^−1^ for Mn).

## Results

### Metagenomic analysis of ANME genomes in Bothnian Sea

Although ANME methanotrophs made up a significant portion of the archaea in the sediments of the two Bothnian Sea sites, retrieving their genomic blueprint appeared challenging. Ultimately, using different assembly and binning parameters as well as differential coverage binning as described in Albertsen et al. (2013), we could recovered eight ANME MAGs from our samples. Two MAGs originated from the sediment of marine Lake Grevelingen (LG) in the Netherlands, and the other six came from either original Bothnian Sea (BS) sediment or the long-term AOM incubations (Table 1).

**Table 1.** ANME MAGs. Genome name as used here, reference, ANME clade, sample origin, number of scaffolds, size, contigs, predicted genes, GC content, Completeness and Contamination based on CheckM2.

| Genome | Reference | Origin | Scaffolds (number) | Size (Mb) | Contigs (number) | Predicted genes (number) | GC (%) | Compl. (%) | Cont. (%) | Strain heterog | Abund. (%) | Seq. method |
| --- | --- | --- | --- | --- | --- | --- | --- | --- | --- | --- | --- | --- |
| GV20_1 | This study | Lake Grevelingen sediment, Netherlands | 118 | 1.9 | 172 | 1994 | 41.8 | 94.4 | 0.7 | 0 | 0.1 | Illumina |
| GV20_2 | This study | Lake Grevelingen sediment, Netherlands | 134 | 1.9 | 191 | 2048 | 41.7 | 98.7 | 0.7 | 0 | 0.1 | Illumina |
| NB8_1 | This study | Bothnian Sea sediment, site NB8 | 141 | 2.1 | 161 | 2475 | 41.8 | 86.7 | 3.9 | 71.43 | 3.1 | Illumina+ Nanopore |
| NB8_2 | This study | Bothnian Sea sediment, site NB8 | 200 | 1.8 | 213 | 1998 | 41.5 | 87.3 | 6.8 | 84.62 | 3.1 | Illumina+ Nanopore |
| US2 | This study | Bothnian Sea sediment, site US2 | 48 | 3.1 | 480 | 3653 | 41.2 | 97.0 | 3.3 | 80 | 2 | Illumina+ Nanopore |
| US2_Fe | This study | Enrichment culture with methane and iron oxides from site US2 | 309 | 1.9 | 401 | 2072 | 42.3 | 90.0 | 0 | 0 | 2.8 | Illumina |
| US5B | This study | Bothnian Sea sediment, site US5B | 679 | 3.1 | 679 | 3590 | 41 | 96.3 | 5.4 | 85.71 | 4 | Illumina |
| US5B_Mn | Klomp et. al. in preparation | Enrichment culture with methane, manganese oxides and molybdate | 281 | 1.7 | 951 | 2119 | 41.7 | 89.5 | 1.3 | 1.31 | 1.8 | Illumina |

**Fig. 1.**
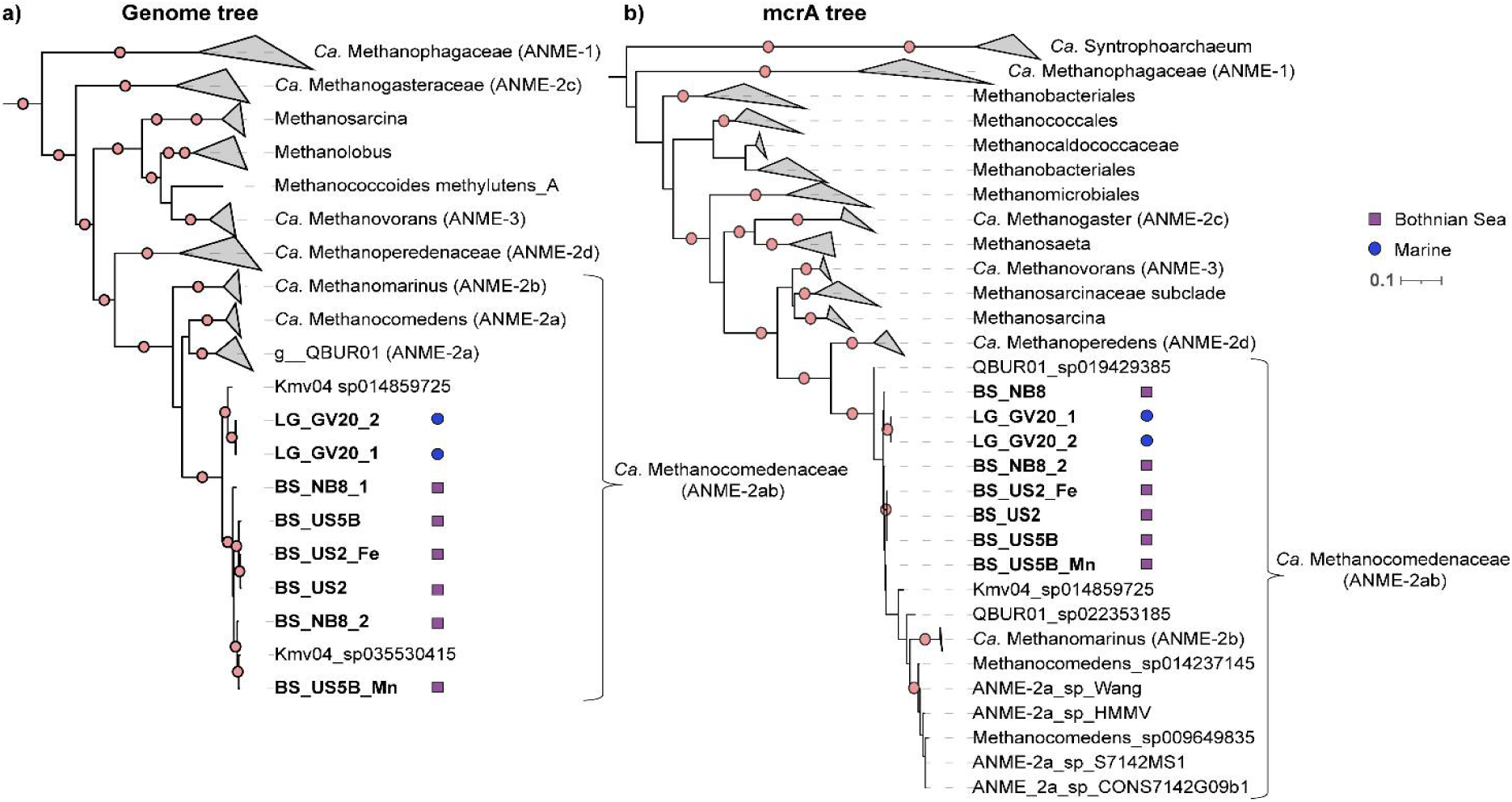
Maximum likelihood phylogenetic tree of a) the assembled genomes based on multiple sequence alignment of 120 concatenated archaeal marker genes and b) *mcrA* sequences of the assembled ANME genomes based on reference file of multiple *mcrA* sequences. Bootstrap support values for 1000 replicates analysis. Dots in the branches indicate 100% bootstrap value. Genomes from this study are in bold and the symbol behind indicate 100% bootstrap value. Genomes from this study are in bold and the symbol behind them indicates if they are from the Bothnian Sea (purple square) or Marine Lake

All MAGs were of high quality, with > 87% completeness and < 7 % contamination. However, all MAGs from the original sediments of the Bothnian Sea sites had very high strain heterogeneity (71-86 %), making it difficult to estimate whether they represent a true individual strain (Table 1). The MAGs also differed in size, from 1.7 Mb (US5B_Mn) to 3.0 Mb (US2 & US5B).

All the recovered ANME MAGs belong to the family *Ca.* Methanocomedenaceae (ANME-2ab) based on multiple sequence alignment of the whole genomes, as well as *mcrA* sequence alignment (Fig. 1). More specifically, all the MAGs belong to ANME-2a genus “Kmv04”, that has only 2 previously published MAGs. With the addition of multiple novel genomes to this ANME-2a cluster, we propose to name this genus “*Candidatus* Methanoborealis” (n. methanum = methane; adj. borealis = northern). Phylogenomically, the BS MAGs cluster closely together with each other and Kmv04_sp035530415, which originates from a soil sample close to the Baltic Sea coast. In comparison, the LG MAGs cluster closely with Kmv04_sp014859725, recovered from a terrestrial mud volcano (Table S3), which is supported by average nucleotide identity (ANI).

The MAGs that originate from the Baltic Sea all share ANI values > 90 %, but have < 90% ANI with the LG MAGs and Kmv04_sp14859725 (Fig. 2). With 95% ANI threshold for species-level identification, all Kmv04 MAGs can be classified as 5 separate species, some with multiple strains within; BS_US2_Fe, BS_US2 and BS_US5B (min. ANI 96.4%); BS_NB8_1; BS_US5B_Mn, BS_NB8_2, Kmv04_sp035530415 (min. ANI 97.1 %); LG_GV20_1 and LG_GV20_2 (min. ANI 99.6 %); and Kmv04_sp011489725. Kmv_04 is also more divergent from the other ANME-2a genera, with average amino acid identity (AAI) < 74.5% to QBUR01 and Methanocomedens, whereas the other two genera share ∼75-77 % AAI based on the selected representative MAGs (Fig. S1). Based on the phylogenomic tree constructed with concatenated marker gene alignment, all the recovered ANME MAGs belong to the family *Ca.* Methanocomedenaceae (ANME-2a; Fig.1).

**Fig. 2.**
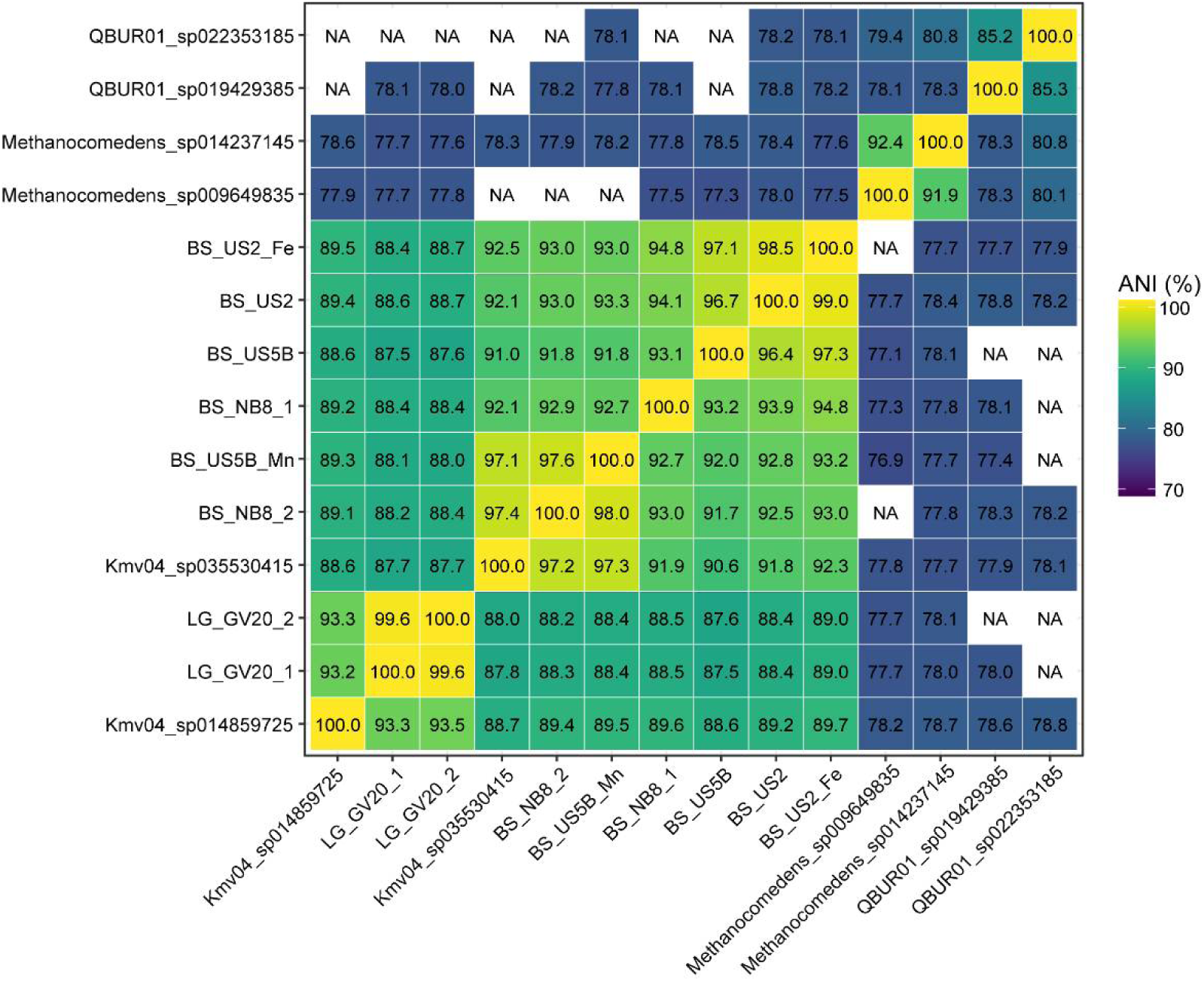
The average nucleotide identity (ANI) between ANME-2a Kmv04 MAGs and other ANME-2a groups; QBUR01 and Methanocomedens. NA = < 70% ANI.

### Metagenomic potential of the ANME-2a Kmv04 cluster

We investigated the relevant functional potential of the BS and LG ANME-2a MAGs and compared them to other ANME-2a genera (Fig. 3). Almost all of our retrieved MAGs encoded the full reverse methanogenic pathway. The analyzed genomes have the potential to fix CO_2_ via the Wood-Ljungdahl pathway (*cdhABCDE)*, although some MAGs miss 1 or 2 of the gens of the pathway. Furthermore, all the MAGs seem to be able to assimilate acetate (*ack*A; Fig. 3; 71 Ouboter *et al.* 2023). Nitrogen fixation potential, via the diagnostic *nif*H from the nitrogenase complex (or other subunit genes; Table S5) were initially only found in the reference Kmv04, and Methanocomedens MAGs, suggesting that nitrogen assimilation may differ between ANME-2a species. However, for NB8_2, we detected a complete cluster of the nitrogenase complex in the metagenome assembly, which had not binned into the MAG.

Genes potentially involved in EET and metal reduction were abundantly present in the BS and LG MAGs. All BS MAGs, encoded *fpoD* from the F420H₂ dehydrogenase complex (Fpo) that is required to transfer electrons to the membrane-crossing electron carriers. NB8_1 and US5B_Mn MAGs were the only genomes where the *rnfC* gene from the membrane-bound ferredoxin:NAD⁺ oxidoreductase complex (Rnf) was mssing. Multiheme OmcZ homologues were detected in the majority of the BS MAGs, as well as in other ANME-2a genera, but not in LG MAGs. All the Bothnian Sea MAGs had collectively a higher number of MHCs than the Kmv04 MAGs from other locations (Fig.3; Table S1). Both LG MAGs only had 10 MHCs, although some of them encoded for large proteins with 42 and 31 heme-binding motifs, respectively. MAG_US5B encoded 52 MHCs, one protein with 70 heme-binding motifs, whereas MAG_US2 had the second highest number with 25 MHCs. Only US5B_Mn did not have any proteins with > 20 heme-binding proteins, although it encoded 15 MHCs in total.

**Fig. 3.**
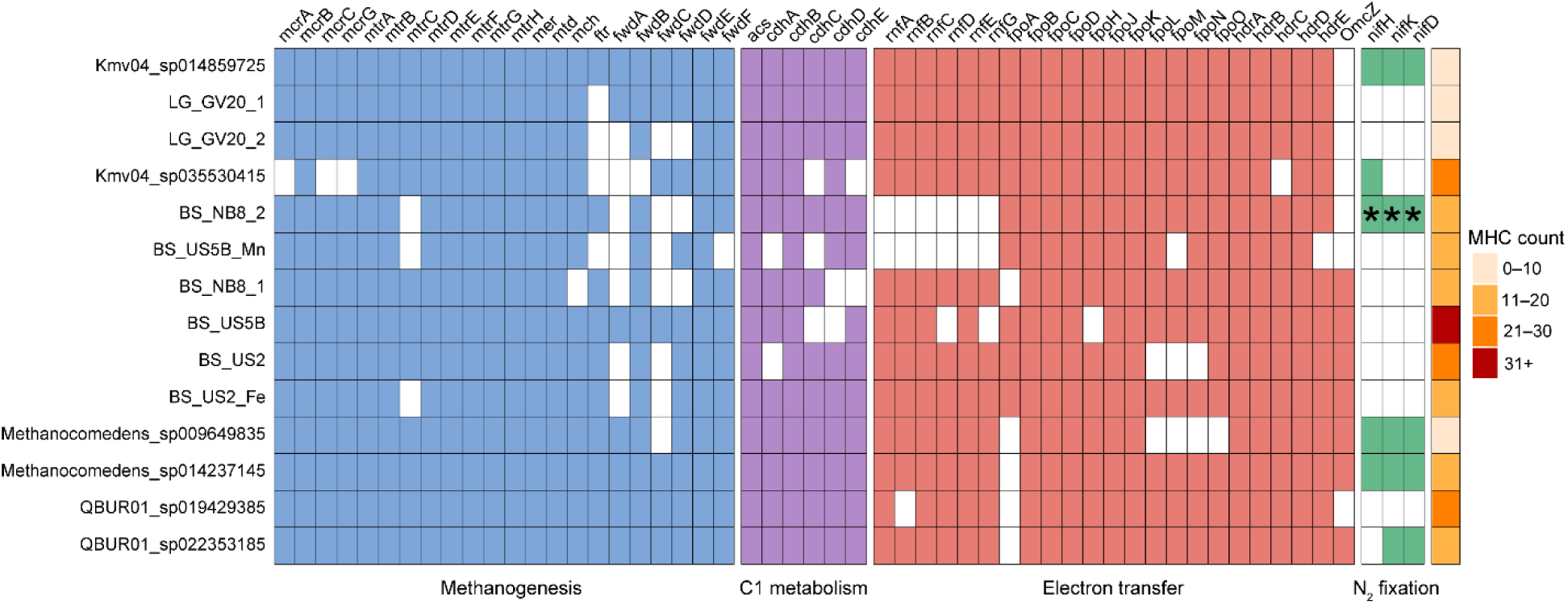
The metabolic potential in ANME-2a genomes for different pathways is shown as a absent/present. Number of multiheme (> 3) cytochrome c proteins is shown on the right column, the colour code representing the range of number of proteins per genome. Marker genes for reverse methanogenesis steps; mcrA-G = Methyl-coenzyme M reductase, mtrA-H = Methyl-H₄MPT:coenzyme M methyltransferase, mer = Methylenetetrahydromethanopterin reductase, mtd = Methylene-H₄MPT dehydrogenase, mch = Methenyl-H₄MPT cyclohydrolase, ftr = Formylmethanofuran:H₄MPT formyltransferase, fwdA-F = Formylmethanofuran dehydrogenase. C1 metabolism genes are in red; acs = Acetyl-CoA synthase, cdhC = CO dehydrogenase subunit C, cdhA = CO dehydrogenase/acetyl-CoA synthase subunit A. Genes for electron transfer and potential metal reduction; rnfA-G = Ferredoxin-NAD⁺ oxidoreductase, fpoA-O = F420H₂ dehydrogenase, hdrA-E = heterodisulfide reductase, OmcZ = Outer membrane c-type cytochrome Z. Nitrogen fixation potential marker gene is in green; nifHKD = nitrogenase iron protein. * indicates a gene retrieved from the metagenome assembly but is not in the MAG.

**Fig. 4.**
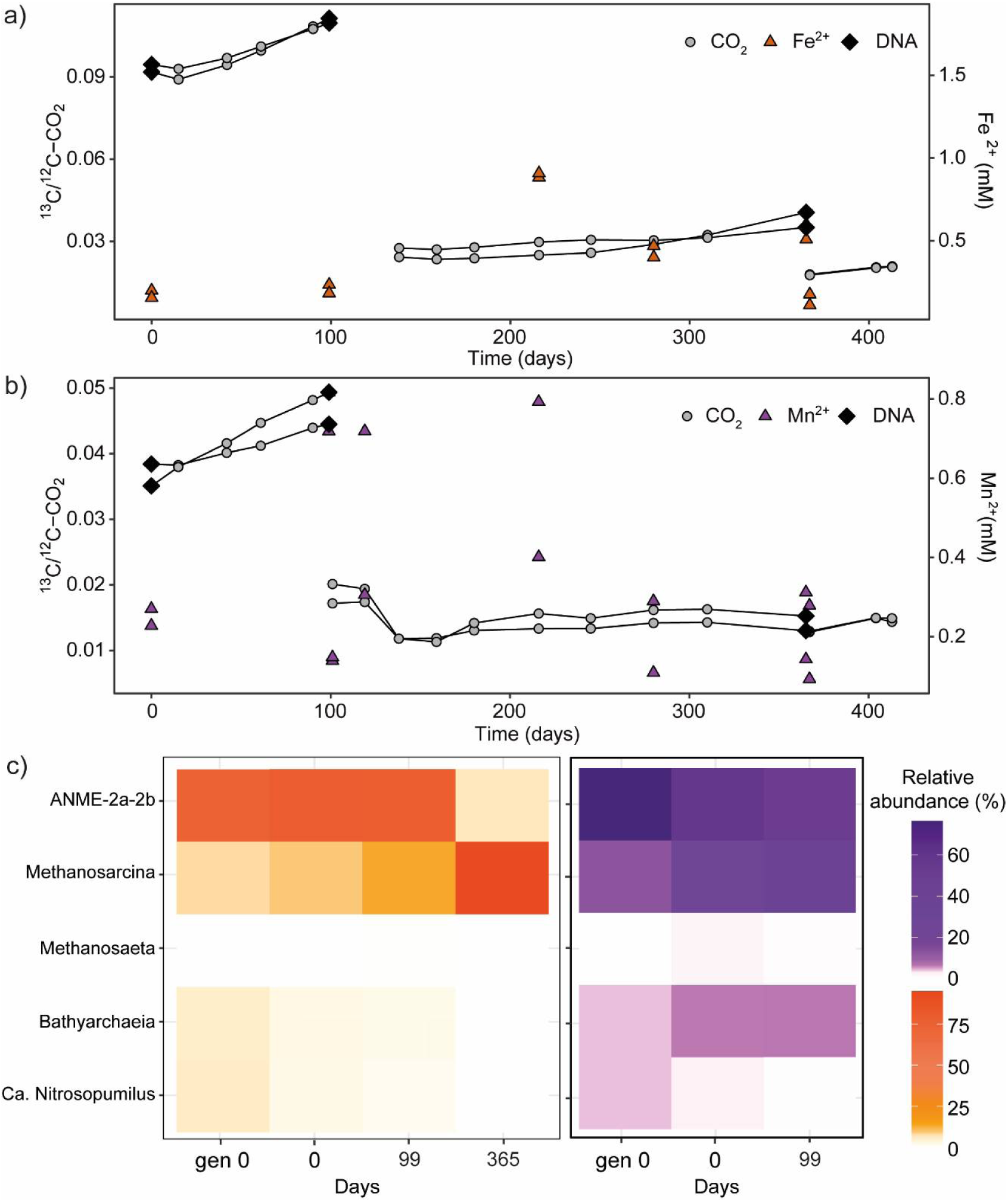
The continuous monitoring of a) Fe-AOM and b) Mn-AOM incubations over 400 days for increase in ^13/12^C-CO_2_ and dissolved metals and c) the relative abundance of the 5 most abundant archaeal genera in the incubations over time based on 16S rRNA amplicon analysis, Fe-AOM on the left (orange/red) and Mn-AOM on the right (purple). The black diamonds represent measurement time points when DNA sample was taken before a 1:5 dilution of the cultures, the dilution also represented as a break in the line.

### Long-term AOM incubations support AOM activity with various electron acceptors

To test if the ANME methanotrophs in the Bothnian Sea sediments can utilize other electron acceptors than sulfate for AOM, as suggested by their genomic potential for metal reduction, we monitored several long-term incubations from two sites in the Bothnian Sea (NB8 & US2). They were amended with ^13^C-labelled methane and either metal oxides or graphene oxide (GO). The samples from US2 amended with manganese (Mn-AOM) and iron (Fe-AOM) showed a clear increase in the ^13/12^C-CO_2_ ratio for the first 100 days after starting new cultures from the original experiment (Fig. 4a-b), indicating active methane oxidation. However, the ^13^CO_2_ production from methane stagnated around day 130 in the Fe-AOM samples following a dilution of 1:5 but slowly re-established around day 250 (Fig. 4a). The concentration of dissolved iron fluctuated, but reached 1 mM on day 220, suggesting active iron reduction activity. In Mn-AOM samples, the signal for ^13^CO_2_ from methane oxidation was not entirely visible after day 100, but this lack of a trend could be explained partly due to the low amount of total headspace CO_2_ and thus ^13^CO_2_ being below detection limit.

We also monitored sulfate concentrations as manganese oxides can oxidize sulfur minerals in the sample. Thus, regardless of using a sulfate-free ASW, sulfate concentrations increased over time, to up to 600 µM on day 100 (Fig. S5). ANME-2ab, according to SILVA taxonomic classification of the 16S rRNA gene amplicon reads, were the most abundant archaea in both incubations for the first 100 days, and the only known methanotroph (Fig. 4c). As metagenome sequencing showed no ANME-2b groups in the samples, we will refer to the population as ANME-2a from now on (Table S2). By day 365, AOM and metal reducing activity together with the relative abundance of ANME had decreased significantly while methanogenic *Methanosarcina* reads now comprised the majority of the archaeal community in Fe-AOM samples (Fig. 4). Due to low biomass and the interference of manganese in DNA extraction, we did not obtain enough DNA for sequencing of Mn-AOM samples on day 365. In the samples with GO, a similar trend of stagnation of AOM was observed between days 100 and 240, but upon re-addition of new electron acceptor the activity was recovered and maintained also after the next dilution on day 365. Likewise, in NOM-AOM samples with GO, ANME-2a was the most abundant archaea with 70% relative abundance on day 0 but a decrease to 20% was observed on day 99 while active methane oxidation was still detected (SFig. 2a-b). It should be noted, however, that the biomass in these samples formed large visible clumps, and possibly attached to the graphene oxide, making it hard to obtain a homogenous sample. Furthermore, graphene oxide adsorbs extracellular nucleic acid during extraction which may have resulted in poor quality DNA, indicated by low number of reads passing quality control during analysis. However, on day 365 the relative abundance of ANME-2a had increased again to 40%, but again we note that only ∼30% of the reads passed to the downstream analysis which could have implications for relative abundances shown here. As observed with Mn-AOM and Fe-AOM samples, *Methanosarcina* reads had the opposite trend to ANME, seemingly taking over the community once ANME and AOM activity decreased and thus *Methanosarcina* was the most abundant archaeal genera on day 365 (Fig. 4). In parallel, we set up enrichments from the shallower site NB8. Here, we also observed Fe-AOM and Mn-AOM for the first 200 days (Fig. S3a-b). With iron, methane oxidation was slow but detectable, with an increase in Fe^2+^. In the manganese incubations, a notable ^13/12^C-CO_2_ ratio increase was not observed after the first dilution, although high dissolved manganese concentrations indicated a reduction of the manganese oxides (Fig. S3b). Again, ANME-2a were the only known methanotrophic archaea in Fe-AOM samples but decreased in abundance over time, while *Methanosarcina* became more abundant (Fig. S3c). In contrast, in the Mn-AOM sample from NB8, ANME-2a relative abundance remained steady for 418 days (∼70 %), while *Methanosarcina* decreased in abundance. Overall, ANME-2a was likely the only microorganism responsible for AOM in all incubations, as their decrease in abundance correlates with a decrease in AOM activity. The presence of multiple strains was also seen in the 16S rRNA gene data, as all incubation samples had 1-2 very abundant Amplicon Sequence Variants (ASVs), and multiple less abundant ones (Table S6). The abundance of the ASVs also varied with different electron acceptors.

In our enrichments, the canonical SRB partners for ANME-2a, *Desulfobacteraceae*, that commonly contains the ANME-2ab-associated SEEP-SRB1 group, decreased in all incubations (Fig. S2, S4, S5). Another common bacterial partner for S-AOM, members of *Desulfobulbales*, did increase or remained at a high abundance in all incubations except with manganese at US2 where they were not detected. In the NB8 samples, *Desulfuromonadaceae* were the most abundant bacteria in both iron and manganese incubations (Fig. S4). Also, *Geobacteraceae* reads increased in the same samples, indicating active metal reduction. Otherwise, the bacterial communities in the Mn-AOM and Fe-AOM samples were very dissimilar.

### Role of *Methanosarcina* in AOM enrichments

As *Methanosarcina* became the dominant archaeal taxa in the Fe-AOM samples of both US2 and NB8 and enriched also in the NOM-AOM samples, we wanted to understand their potential role in the metal- and methane cycling in our samples. We recovered one *Methanosarcina spp.* MAG from each site in NB8 and US2 and investigated their metabolic potential (Fig. 5). The MAGs were ∼78 % complete, which might explain why they did not encode *mcrA*, *mch* or *ftr*; all necessary for methanogenesis (Table S8). However, *mcrB* was present in both MAGs. Also genes from acetoclastic (*ack*A, *pta*) and methylotrophic (*mtaB, mttB, mtbB*) methanogenesis were present (Fig. 5). Both MAGs had most subunits for the electron transfer complexes Rnf, Fpo and Hdr. Methanosarcina_NB8 encoded 4 c-type cytochromes with 3 or five heme-binding motifs, where Methanosarcina_US2 had 3 cytochrom c. One 5-heme c-type cytochrome in the US2 genome was identified as outer-surface multiheme c-type cytochrome MmcA (Table S9).

**Fig. 5.**
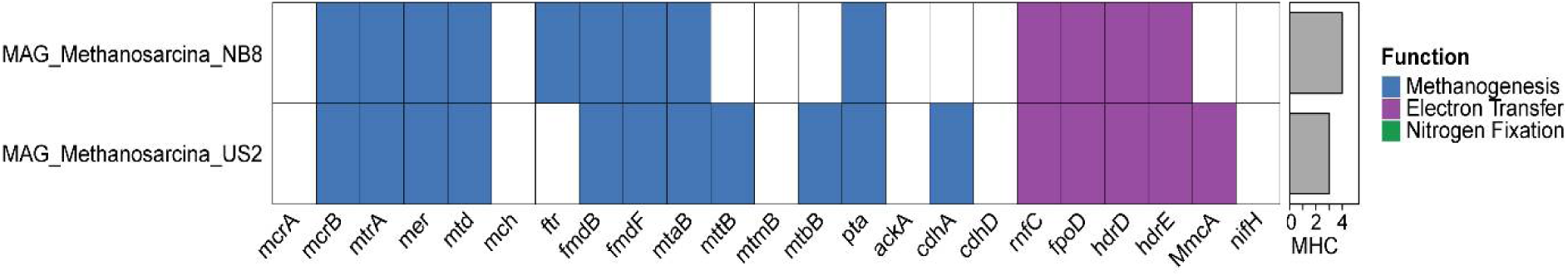
Metabolic potential of *Methanosarcina* MAGs. The presence or absence of gene of different metabolic pathways. Number of multiheme (> 3) cytochrome c proteins is shown on the right bar plot. Marker genes for reverse methanogenesis steps are shown in blue; genes for electron transfer and potential metal reduction are in purple; nitrogen fixation potential marker gene is in green (not present). Protein annotations for both MAGs from DRAM are in Table S8.

## Discussion

### Novel ANME-2a genus with metal-AOM potential

Recovering high-quality genomes from complex sediment samples remains a significant challenge due to low abundance and high strain diversity [55]. In our study, we addressed these challenges by applying both short and long read sequencing and a combination of assembly and binning strategies, including differential coverage binning (Albertsen et al., 2013), which allowed us to reconstruct eight high-quality ANME-2a MAGs from the Bothnian Sea and Lake Grevelingen sediments. Despite high completeness and low contamination, most MAGs from Bothnian Sea sediments displayed high strain heterogeneity, making strain characterization complicated in the presence of many closely related strains, a common issue in metagenome-resolved population genomics [56].

Phylogenomic analyses based on concatenated marker genes and *mcrA* sequences consistently placed all MAGs within the ANME-2a clade of the family *Ca.* Methanocomedenaceae [7]. Phylogenomically closest to our MAGs are two genomes assigned to the clade provisionally named “Kmv04”. These 2 genomes are so far the only representatives for this lineage, despite originating from geographically and environmentally distant sites (>4000 km apart). The observed AAI values (60–80%) fall within the range typically associated with genus-level separation (Rodríguez-R and Konstantinidis, 2014), supporting the view that these genomes represent a distinct taxonomic group. [57,58]. Based on the available evidence, we therefore propose to name this new cluster within ANME-2a as genus “*Candidatus* Methanoborealis” (n. methanum = methane; adj. borealis = northern).

As indicated by the ANI values, the different clusters suggest distinct sub lineages within *Ca.* Methanoborealis adapted to different environments, and possibly habitat-specific adaptations which are common for ANME [59]. Interestingly, the habitat of the GV MAGs is much more different from the mud volcano sediments than the Bothnian Sea, suggesting a special Bothnian Sea niche that has evolved separately from the other *Ca.* Methanoborealis species. However, more MAGs representing this genus from different ecosystems are needed to validate the niche separation. Within the BS subgroup, the species are closely related and some, just as US5B, US2 and US2_Fe seem to represent different strains of the same species with ANI > 95% [60].

The separate clusters within the genus were also apparent in the metabolic potential of the MAGs. The *Ca.* Methanoborealis MAGs showed notable differences in metabolic potential, especially regarding EET and metal reduction capacities. The BS MAGs encoded a substantially higher number of MHCs, including large proteins with more than 70 heme-binding motifs. The presence of large MHCs has been linked to capacity to transfer electrons to extracellular electron acceptors such as metal oxides [2,10]. This suggests a stronger genomic potential for metal-dependent AOM in the BS lineage, particularly in the US2 and US5B MAGs. Both sites are characterized by burial of metal oxides below the SMTZ, and at US5B, ANME-2ab is highly abundant below this zone [15,61]. The presence of OmcZ homologue in most of the BS MAGs also supports metal-AOM potential, as in *Geobacter* this MHC is required for nano-wire formation in EET [25]. However, OmcZ is commonly found in ANME-2ab genomes, as also seen in the QBUR01 and *Methanocomedens* genomes, but its exact function or role in EET is not known [7]. *Ca.* Methanoperedens species often have several large MHCs whose proposed function is EET. However, the specific MHCs used by ANME have not been identified and the proteins used in the presence of extracellular electron acceptors appear to differ depending on the substrate and species, based on gene expression studies[4,10,12,62].

IIn Lake Grevelingen sediments, ANME-2ab have been only found to be active around the SMTZ, suggesting that S-AOM is their main metabolism [27], which could explain the lower number and smaller MHCs encoded and the lack of OmcZ in their genomes. Notably, nitrogen fixation genes such as nifH were absent or not binned in most of our MAGs, but present in the reference Kmv04 MAG and in *Methanocomedens*, aligning with previous work suggesting that not all ANME-2a populations retain a diazotrophic potential [7].

These metabolic differences, together with the ANI values and phylogenetic clustering, support the conclusion that *Ca.* Methanoborealis includes ecophysiologically diverse populations adapted to varying geochemical conditions, where the BS cluster seems to have a high potential for metal-AOM.

### ANME-2a maintain methane oxidizing activity with various electron acceptors

In addition, we studied the AOM pathways and microorganisms with batch incubations using brackish, methane- and metal-oxide rich coastal sediments. As sulfate concentrations are low in the methanic zone, systems like the Bothnian Sea are potential hotspots for metal-AOM [63,64]. Fe-AOM activity, as well as a high abundance of ANME-2ab in the metal-oxide rich zone below the SMTZ in Bothnian Sea sediments, has been observed before [15,65]. Here, we aimed to better understand their wider occurrence across sites and use of electron acceptors, and to characterize the metabolic pathways involved in metal-AOM with long-term incubations and metagenomic analysis. In all AOM incubations from two Bothnian Sea sites, US2 and NB8, ANME-2a were the only canonical archaeal methanotrophs present. In US2 samples, we detected AOM activity with all added electron acceptors (Mn-oxides, Fe-oxides and graphene oxide), and ANME-2a covered > 60% of total archaeal reads, indicating them to be most likely responsible for methane oxidation. Similarly, in the NB8 samples, amended with Fe- and Mn-oxides, ANME-2a were the most abundant archaea and the only archaeal methanotrophs, and there was a clear ^13^C-enriched signal for AOM, especially with Fe-oxides. Thus, together these results are very strong indications that ANME-2a are the most likely candidates for metal-AOM in these sediments. ANME-2a have been reported to be abundant in the Baltic Sea sediment before [13–15,66], which supports our finding that they are significant players in the Bothnian Sea methane cycle. However, due to the presence of sulfate even after 600 days of incubation, we cannot exclude a cryptic sulfur cycle. Especially in Mn-AOM incubations, Mn oxides might oxidise solid sulfur compounds and thus release sulfate into the medium to drive this cryptic sulfur cycling [67].

Although ANME-2a were the dominant methanotrophs during the initial active phase in US2 Fe-AOM incubations (0–100 days), and for > 400 days in NB8 samples, their relative abundance declined over time, particularly after we diluted the enrichments several times. Instead, *Methanosarcina* appeared to take over the archaeal community in iron and GO amended samples. Although many *Methanosarcina* strains were present, we could retrieve and analyse two *Methanosarcina* MAGs, one from US2 and one from NB8 (Table S7). Due to the medium-quality of the MAGs, we were not able to recover all methanogenesis pathway genes in the genomes, making the analysis cumbersome. We searched for MHCs in the MAGs, as *Methanosarcina* has been shown to be able to reduce iron via EET that involved c-type cytochromes both during acetoclastic and methylotrophic methanogenesis (Guo and Lu, 2025; Holmes et al., 2019). This process is often pronounced in the presence of electron carriers such as humic acid analogue AQDS. *Methanosarcina* has even been shown to be able to reverse methanogenesis and perform Fe-AOM (Yu et al., 2022). We found most components of membrane-bound electron transfer complexes Rnf, Fpo and Hdr in both MAGs, indicating that they belong to Type II methanogens that are capable of EET via MHCs [68]. The US2 MAG encoded MmcA membrane-bound multiheme c-type cytochrome that is upregulated during iron oxidation in *Methanosarcina* and the key component in EET [69], confirming that the *Methanosarcina* in our enrichments are capable of iron reduction. Therefore, we propose that in our cultures, ANME might be outcompeted by *Methanosarcina* for iron oxides and GO, used either for methanogenesis with remaining acetate and methylated substrates in the samples, or Fe-AOM. Since ANME are notoriously slow growers, over several dilutions of the sample, their biomass could have been too low to establish a stable community and they may thus have been outcompeted by *Methanosarcina*. Additionally, both ANME and *Methanosarcina* were potentially active in the incubations, but the weak AOM signal could be masked partly by increased methane production. In US2 Mn oxide incubations, the *Methanosarcina* increase was a lot slower, and in NB8 samples, *Methanosarcina* reads decreased over time, indicating that different competitive metabolism processes occurred. Regardless, our results demonstrate that ANME-2a is capable of metal-AOM in long-term incubations, but their slow growth and unknown metabolic preferences make continuous culturing challenging.

Interestingly, across all incubations, canonical sulfate-reducing bacterial (SRB) partners of ANME-2a —particularly *Desulfobacteraceae* (e.g., SEEP-SRB1)—were either undetectable or decreased over time. This suggests that the ANME-2a lineages enriched here are capable of metal reduction independent of syntrophic SRB, aligning with genomic predictions and prior reports of sulfate-independent ANME metabolism [23,70]. Alternatively, the syntrophic partner might be a metal-reducing partner, such as *Desulfobulbaceae*, as suggested for ANME-2a in Fe-AOM [24]. Indeed, bacterial taxa known for metal reduction, such as Desulfobulbus, *Desulfuromonadaceae* and *Geobacteraceae*, were prevalent in several iron and manganese incubations, implying a potential ecological interaction or competition with ANME for electron acceptors.

## Conclusion

Here, we investigated the metal-AOM potential of ANME-2a clades in metal- and methane rich sediments. With the recovery of 8 novel ANME-2a MAGs, we identified a novel genus of ANME-2a for which we propose a name *Ca.* Methanoborealis. Six of the MAGs were recovered from the sulfate-free layers and metal-oxide rich sediments of the Bothnian Sea, and showed high potential for metal-AOM in their genomes. The other two originate from sulfate-rich zones of marine Lake Grevelingen and showed less potential for metal-AOM, based on the low number of multi-heme cytochrome c type proteins and the absence of OmcZ-like protein. The long-term AOM incubations showed a persistent community of ANME-2a that coincided with methane oxidation and metal reduction activity. However, in the long term, versatile *Methanosarcina* seemed to replace ANME as the dominant archaea, likely outcompeting them for metal oxides and graphene oxide. Together, our data shows that the sublineage of ANME-2a, *Ca.* Methanoborealis have a high diversity in their metabolic pathways and at least the Bothnian Sea subgroup is likely to be an important player in the methane cycle via metal-dependent AOM. The addition of multiple novel ANME MAGs from an under-represented lineage contributes to our growing understanding of the diversity and metabolic flexibility of ANME archaea and highlights how much we still have to learn about the fascinating and important process of metal-dependent-AOM.

## Acknowledgements

This research was funded by NESSC NWO-OCW 024002001, SIAM NWO-OCW 024002002 440 and ERC MARIX 854088. We thank all MARIX members for assistance in sampling and acquiring of material for the MAGs from in-situ sediments.

## Data availability

All sequencing data originating from this and previous studies have been uploaded to the National Center for Biotechnology Information (NCBI) website under the following BioProject accession numbers: PRJNA511814 US5B Bothnian Sea 2019; PRJNA1310174 and PRJNA1167897 Scharendijke 2020, Lake Grevelingen NL. The raw sequencing data of US5B 2023 have accession number PRJNA1314788. The data for site US2 are deposited under BioProject accession number PRJNA1310174 and PRJNA1306556. All other data of US5B 2023 are deposited on Zenodo at 10.5281/zenodo.17174800. All other data for this study are uploaded to Zenodo data repository per 9/1/26 under https://doi.org/10.5281/zenodo.16937456.

