## Supplementary material for "Potential for metal-coupled methane oxidation by *Candidatus* Methanocomedenaceae in coastal sediments": suppl fig

### Supplementary Material Methanoborealis ms Wallenius et al 2026

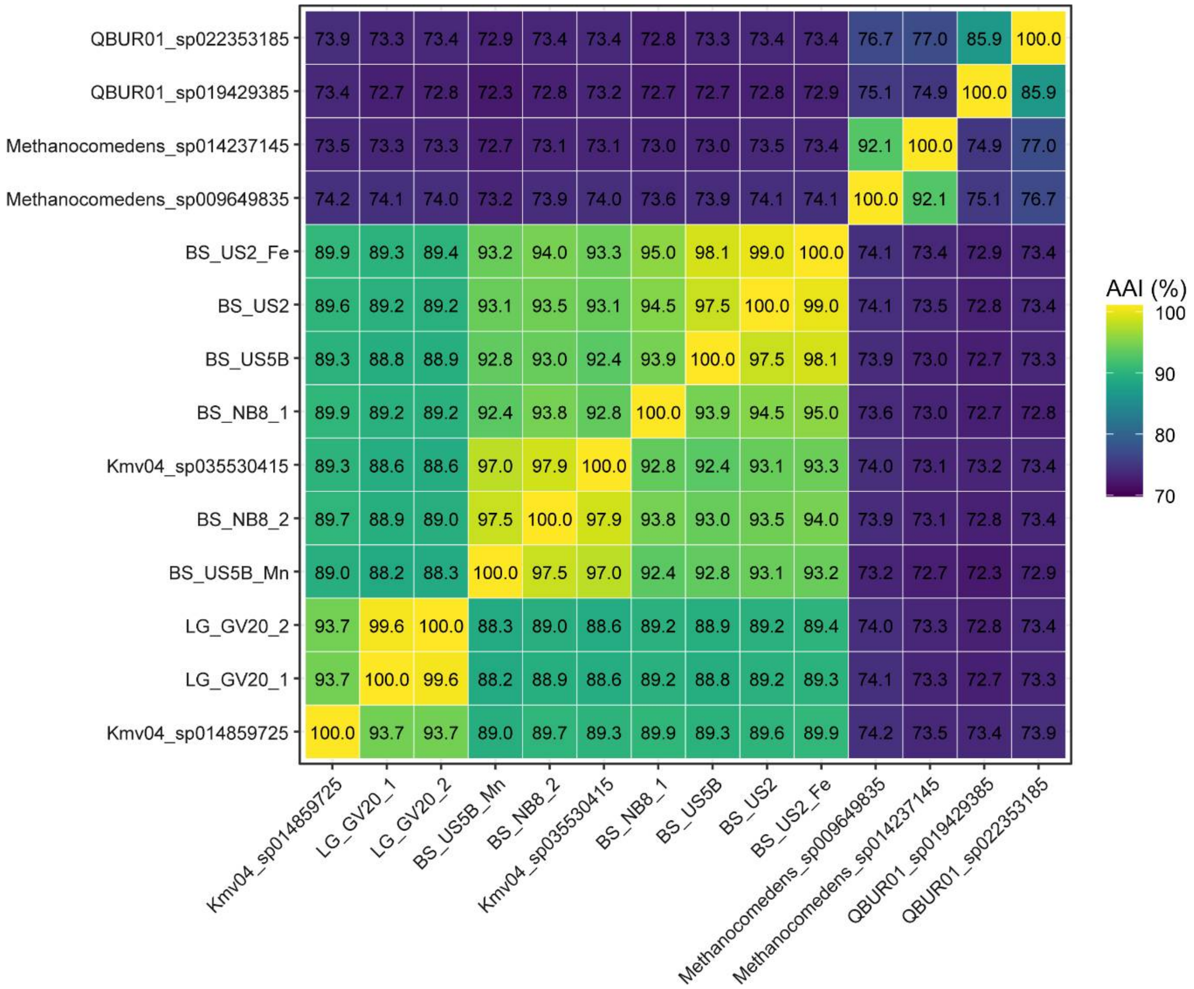


**Fig. S1.** The average amino acid identity (AAI) between ANME-2a Kmv04 MAGs and other ANME-2a groups; QBUR01 and Methanocomedens.

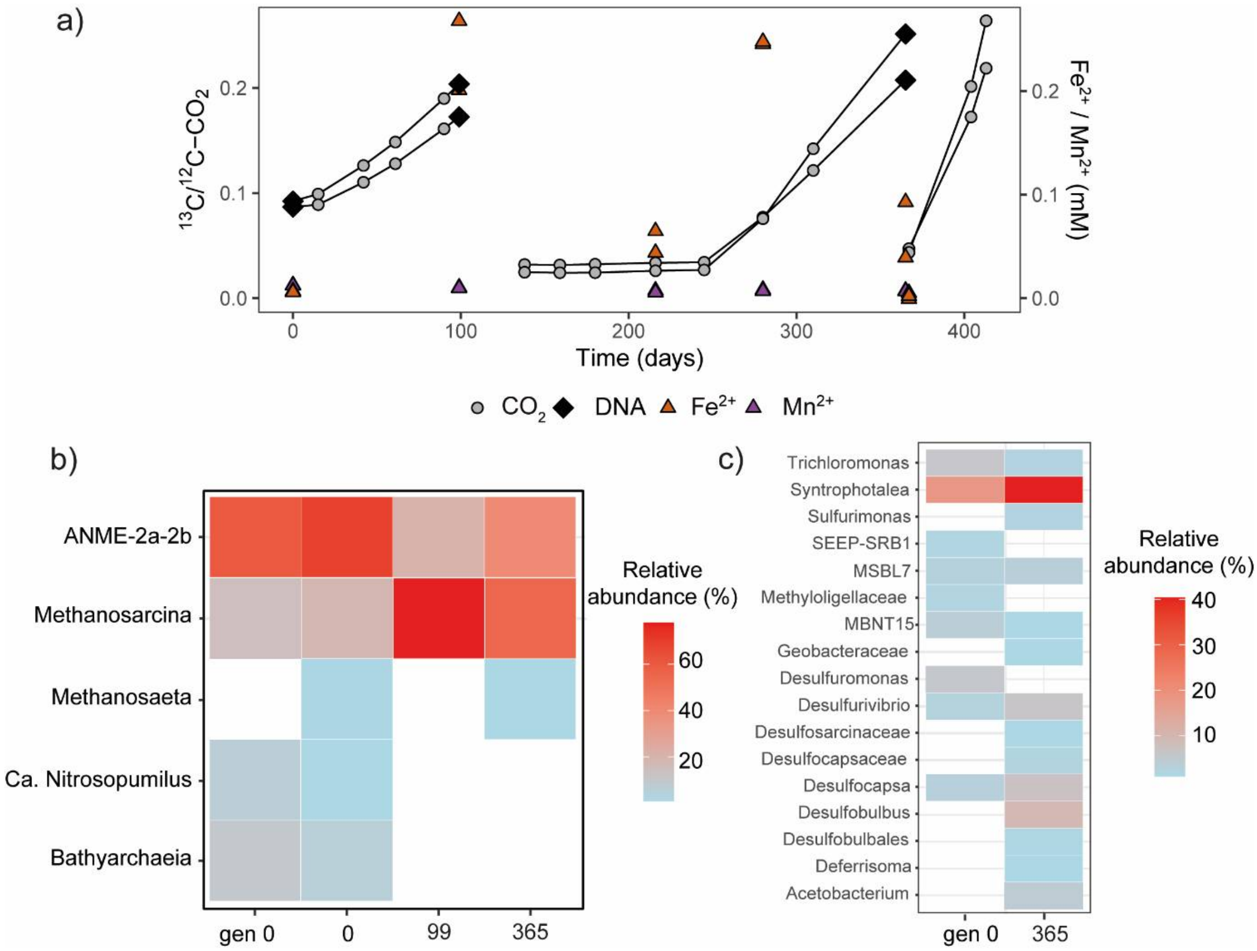


**Fig. S2.** The methane oxidation activity as $^{13/12}CO_2$ ratio in graphene oxide amended samples from US2 and concentration of dissolved iron and manganese in the medium (a) and the relative abundance of top archaeal (b) and bacterial (c) taxa over time based on 16S rRNA gene amplicon sequencing. The black diamonds represent measurement time points when a DNA sample was taken before a 1:5 dilution of the cultures, the dilution is also represented as a break in the line.

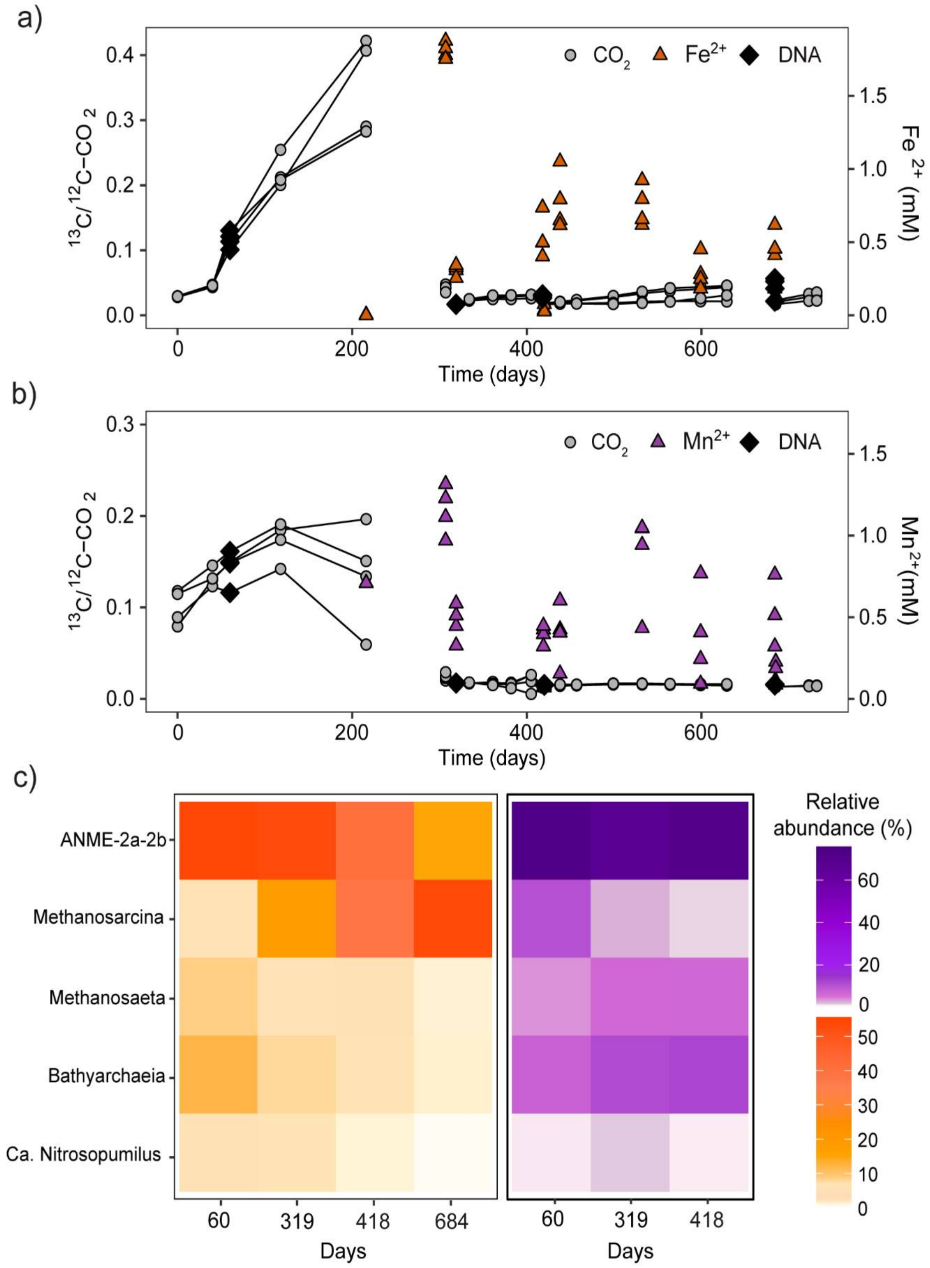


211

**Fig. S3.** The methane oxidation activity as $^{13/12}CO_2$ ratio and dissolved metal concentration in a) iron oxide and b) manganese oxide-amended samples from NB8 and c) the relative abundance of top archaeal taxa over time based on 16S rRNA gene amplicon sequencing. Left/orange = Fe-AOM; Right/purple = Mn-AOM. The black diamonds represent measurement time points when DNA sample was taken before a 1

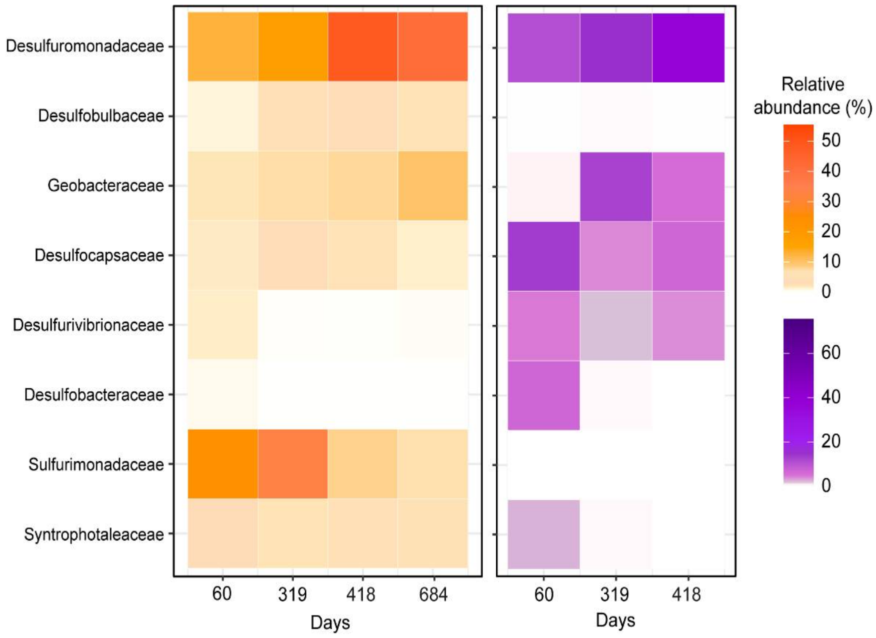


**Fig. S4.** Relative abundance of selected bacterial families in NB8 Fe-AOM (a) Mn-AOM (b) incubations over time. 0 = 405 days; 1 = 665 days; 2 = 765 days and 3 = 1030 days.

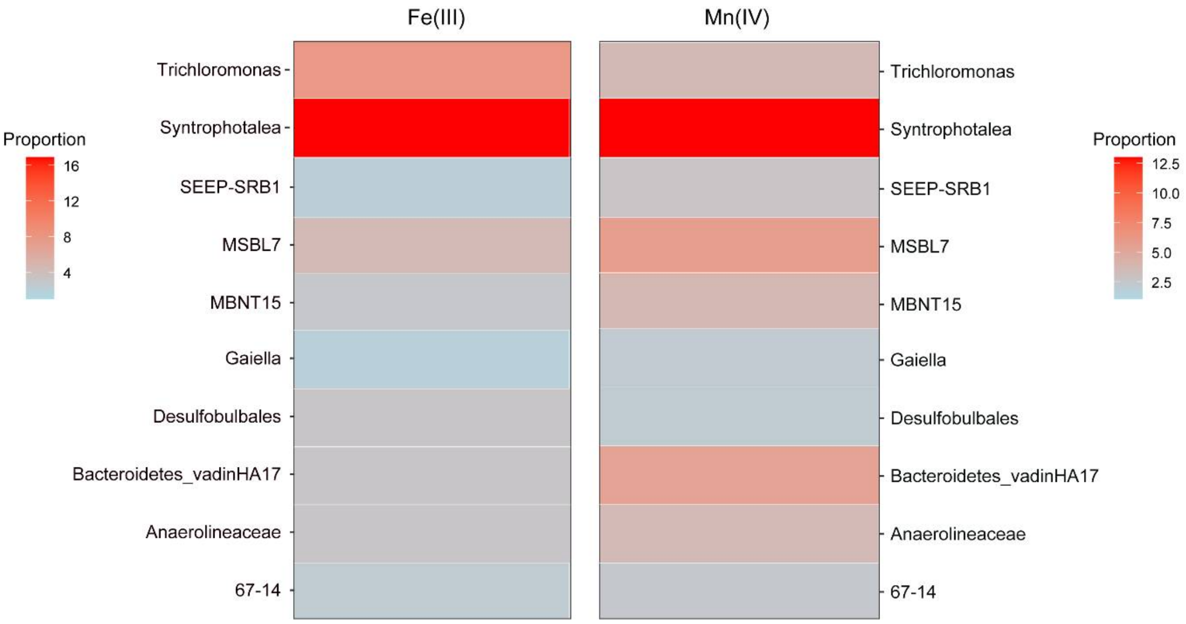


**Fig. S5.** Relative abundance of selected bacterial families in US2 Fe-AOM (a) Mn-AOM (b) incubations before the beginning of this study (gen0).

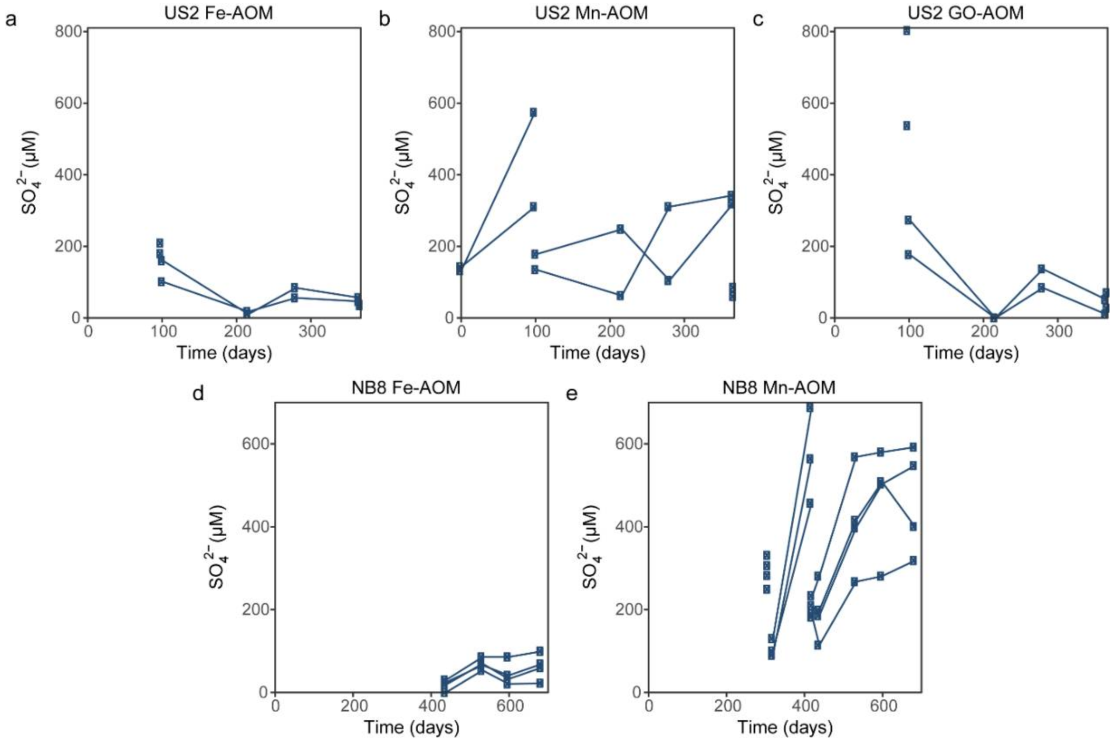


**Fig. S6.** Sulfate concentrations in the US2 incubations (a-c) and NB8 (d-e) over time in all separate samples, with replicates shown in their own lines, over time. Breaks in lines represent dilution of samples when sulfate-free ASW was used in a 1:5 dilution.

**Table S1.** The amount of MHC proteins in Kmv04 and Methanosarcina genomes and the maximum number of heme-binding motifs in a single protein.

| **Genome** | **Number of MHC** | **Max. heme-binding groups** |
|---|---|---|
| Kmv04_sp014859725 | 10 | 34 |
| Kmv04_sp035530415 | 22 | 43 |
| MAG_GV20_1 | 10 | 42 |
| MAG_GV20_2 | 10 | 31 |
| MAG_NB8_1 | 18 | 34 |
| MAG_NB8_2 | 14 | 34 |
| MAG_US2 | 25 | 51 |
| MAG_US2_Fe | 12 | 51 |
| MAG_US5B | 52 | 70 |
| MAG_US5B_Mn | 15 | 16 |
| MAG_Methanosarcina_NB8 | 4 | 5 |
| MAG_Methanosarcina_US2 | 3 | 5 |

**Table S2.** All Methanocomedenaceae reads and their coverage in Fe-AOM incubation metagenomes based on singleM analysis.

| Sample | Coverage (Root) | Taxonomy (GTDB) |
|---|---|---|
| NB8-Fe-gen0 | 2 | d__Archaea; p__Halobacteriota; c__Methanosarcinia; o__Methanosarcinales; f__Methanocomedenaceae |
| NB8-Fe-gen0 | 57.68 | d__Archaea; p__Halobacteriota; c__Methanosarcinia; o__Methanosarcinales; f__Methanocomedenaceae; g__Kmv04 |
| NB8-Fe-gen0 | 0.44 | d__Archaea; p__Halobacteriota; c__Methanosarcinia; o__Methanosarcinales; f__Methanocomedenaceae; g__QBUR01 |
| NB8-Fe-gen0 | 21.41 | d__Archaea; p__Halobacteriota; c__Methanosarcinia; o__Methanosarcinales; f__Methanocomedenaceae; g__Kmv04; s__Kmv04 sp014859725 |
| NB8-Fe-1 | 2.44 | d__Archaea; p__Halobacteriota; c__Methanosarcinia; o__Methanosarcinales; f__Methanocomedenaceae |
| NB8-Fe-1 | 41.62 | d__Archaea; p__Halobacteriota; c__Methanosarcinia; |

| | | |
|---|---|---|
| | | o__Methanosarcinales; f__Methanocomedenaceae; g__Kmv04 |
| NB8-Fe-1 | 14.63 | d__Archaea; p__Halobacteriota; c__Methanosarcinia; o__Methanosarcinales; f__Methanocomedenaceae; g__Kmv04; s__Kmv04 sp014859725 |
| NB8-Fe-4 | 2.91 | d__Archaea; p__Halobacteriota; c__Methanosarcinia; o__Methanosarcinales; f__Methanocomedenaceae |
| NB8-Fe-4 | 30.28 | d__Archaea; p__Halobacteriota; c__Methanosarcinia; o__Methanosarcinales; f__Methanocomedenaceae; g__Kmv04 |
| NB8-Fe-4 | 0.64 | d__Archaea; p__Halobacteriota; c__Methanosarcinia; o__Methanosarcinales; f__Methanocomedenaceae; g__QBUR01 |
| NB8-Fe-4 | 12.66 | d__Archaea; p__Halobacteriota; c__Methanosarcinia; o__Methanosarcinales; f__Methanocomedenaceae; g__Kmv04; s__Kmv04 sp014859725 |
| US2-Fe-1 | 1.85 | d__Archaea; p__Halobacteriota; c__Methanosarcinia; o__Methanosarcinales; f__Methanocomedenaceae |
| US2-Fe-1 | 70.66 | d__Archaea; p__Halobacteriota; c__Methanosarcinia; o__Methanosarcinales; f__Methanocomedenaceae; g__Kmv04 |
| US2-Fe-1 | 29.62 | d__Archaea; p__Halobacteriota; c__Methanosarcinia; o__Methanosarcinales; f__Methanocomedenaceae; g__Kmv04; s__Kmv04 sp014859725 |

**Supplementary Tables S3-S9** are available in Zenodo Data repository at https://doi.org/10.5281/zenodo.16937456

Access link:

https://zenodo.org/records/16937456?token=eyJhbGciOiJIUzUxMiJ9.eyJpZCI6ImFjNGQyM2EwLTY5MjUtNGVlYS1hMjQ4LTJhMzhiODg5MjliZCIsImRhdGEiOnt9LCJyYW5kb20iOiI2MmY2MDY5Yzk4ZmMwZTgxZmJkMjljMjg2MGYzZDJhMiJ9.TOxpT3X0SztDb0ewwvXQ-WCWko2OiauBv3s3eAr7JNBLo1TCs_vgCodOMbEF8_nzPqmEBeqz3gl-4LCvPO1QMw

#### Supplementary methods

##### ASW composition

ASW NB8: following composition per L: 3.5 g NaCl, 0.22 g $CaCl_2 \cdot 2H_2O$, 0.1 g KCl, 0.012g $MgSO_4 \cdot 7H_2O$, 0.05g $KH_2PO_4$, 0.05 $NH_4Cl$ and 200 uL of trace element solution prepared as described before [8] and the pH was adjusted to 7.5 with 1 M NaOH.

ASW US2: 5.2 g NaCl, 0.28 g $CaCl_2 \cdot 2H_2O$, 0.1 g KCl, 0.02g $MgCl_2 \cdot 7H_2O$, 0.05g $KH_2PO_4$, 0.05 $NH_4Cl$ and 200 uL of trace element solution prepared as described before [8] and the pH was adjusted to 7.5 with 1 M NaOH.

##### Recovery of US5B MAGs

Site US5B is located in the deepest basin in the Bothnian Sea and is characterized by high porewater concentrations of $Fe^{2+}$ and $Mn^{2+}$ and potential for iron-mediated AOM [3]. The sediments were collected during a research cruise with the R/V Pelagia in July 2023. To recover the BS_US5B MAG, the DNA was extracted from US5B sediments (Table S1) and sequenced with Illumina as described in the methods section. The quality control, assembly and binning was done as follows:

BBDuk (BBTools package v37.76; Bushnell et al., 2014) was used for quality trimming and filtering of the raw sequencing reads (adapter trimming using the BBDuk-supplied adapters.fa reference with k=23, mink=11, hdist=1, tpe, tbo; quality trimming and filtering using qtrim=r, ftm=5). Tadpole (BBTools v39.06) was used for error correction (mode=correct, k=50) and BBnorm (BBTools v37.76) was applied to normalize the coverage of the reads (target=30, min=2). Coassembly of the trimmed and filtered reads was conducted by metaSPAdes (v3.15.5; Nurk et al., 2017) with k-mer sizes 21, 33, 55, 77, 99, 121 and using the --only-assembler flag. Per sample, the reads were mapped back to the assembled contigs with BBMap (BBTools v37.76; slow=t), and the mapping files were processed using SAMtools (v1.19.2; Li et al., 2009).

Binning of the coassembled contigs was done with CONCOCT (v1.1.0; Alneberg et al., 2014), MaxBin (v2.2.7; Wu et al., 2016), MetaBAT 2 (v2:2.15; Kang et al., 2019), and SemiBin2 (v2.0.2; Pan et al., 2023). Consensus bins were generated by DAS Tool (v1.1.2; Sieber et al., 2018). The quality of the consensus bins was checked with CheckM2 (v1.1.2; Chklovski et al., 2023), and GTDB-Tk v2 (v2.4.0; Chaumeil et al., 2022) was used for the taxonomic classification of the bins. have been uploaded to the National Center for Biotechnology Information (NCBI) website under the BioProject accession number PRJNA1283585.

The BS_US5B_Mn MAG was recovered from sediment incubations amended with birnessite, molybdate and 13C-labelled CH4 after 178 days of incubation. Briefly, the sediment was diluted with sterile sulfate free ASWin a 1:4 ratio under a $N_2$ atmosphere for a final amount of 60 g in 120 ml serum bottle and the electron acceptors were added before the bottles were capped and sealed. The headspace was replaced by a mixture of 76% N2, 4% CO2 and 20% 13C-labelled CH4.. and the samples were incubated on a shaking table (90 rpm) at 4°C in the dark. The DNA extraction, metagenomic sequencing, assembly and binning was done as described in the main text for US2 and NB8 samples.

**Recovery of GV MAGs**

The Lake Grevelingen in the Netherlands suffers from eutrophication-driven deoxygenation and euxinia in the bottom waters. This has led to high methane efflux from the sediments due to dysfunctional methane-filter [4,5]. The sediment samples were obtained from Scharendijke basin, the deepest point of the Lake, R/V Navicula in September 2020 [6]. The GV MAGs were recovered from original sediment samples within the sulfate-methane transition zone (Table S1), while the DNA extraction, metagenomic sequencing, assembly and binning was done as described in the main text.

**Table S3**. Sample site description and SRA ID for samples where ANME MAGs in this study originated.

| MAG | Type | Location | Depth water column (m) | Depth Sample (cmbsf) | Sampling time | BioProject Raw reads |
|---|---|---|---|---|---|---|
| GV20_1 | In-situ sediment | 51.742°N, 3.849°E | 45 | 9-11 | September 2020 | PRJNA1167897 SRR33663671 |
| GV20_2 | In-situ sediment | 51.742°N, 3.849°E | 45 | 15-17 | September 2020 | PRJNA1167897 SRR33663672 |
| NB8_1 | In-situ sediment | 63°29.1'N, 19°49.5'E | 34 | 33-36 | June 2019 | PRJNA1309620 *not submitted |
| NB8_2 | In-situ sediment | 63°29.1'N, 19°49.5'E | 34 | 50-53 | June 2019 | PRJNA1309620 *not submitted |
| US2 | In-situ sediment | 62˚50.99' N; 18˚53.53' E | 202 | 28-32 | May 2022 | PRJNA1306556 |
| US2_Fe | $Fe_{ox}$ amended sediment incubation | 62˚50.99' N; 18˚53.53' E | 202 | 28-32 | May 2022 | PRJNA1310174 *not submitted |
| US5B | In-situ sediment | 62.5862°N, 19.9688°E | 214 | Top 300 m (co-assembly) | July 2023 | PRJNA1283585 |
| US5B_Mn | $Mn_{ox}$ amended sediment incubation | 62.5862°N, 19.9688°E | 214 | 75.75 – 78.75 & 95.75 – 98.75 | July 2023 | PRJNA1314788 |
